# *Lasiodiplodia* species associated with mango (*Mangifera indica* L.) decline in Burkina Faso and influence of climatic factors on the disease distribution

**DOI:** 10.1101/2023.03.25.533858

**Authors:** Oumarou Zoéyandé Dianda, Issa Wonni, Léonard Ouédraogo, Philippe Sankara, Charlotte Tollenaere, Emerson M. Del Ponte, Diana Fernandez

## Abstract

Dieback or decline caused by *Lasiodiplodia* spp. is a major disease of mango trees (*Mangifera indica* L.). The main objectives of this study were to identify *Lasiodiplodia* species associated with mango decline in Burkina Faso, and to asses the climatic and edaphic factors affecting the geographic distribution of the disease in the country. The genetic diversity of 47 *Lasiodiplodia* isolates was studied based on sequence data of the translation elongation factor 1-alpha gene (*tef1-a*) and the rDNA internal transcribed spacer region (ITS). Phylogeny analyses grouped the isolates from Burkina Faso with 6 different *Lasiodiplodia* species, including *L. euphorbicola* that accounted for 36 of the 47 isolates. *Lasiodiplodia* isolates tested on mango seedlings induced the typical dieback symptoms. Disease incidence and severity were generally higher in the drier and warmer regions (eastern) of the country. This study provides the required information to establish control strategies against mango decline in Burkina Faso.

## 1. Introduction

Mango (*Mangifera indica* L.) produces delicious fruits and is one of the most important fruit crops in the world’s tropical and subtropical areas. In Burkina Faso, mango occupies the first place in fruit production, and marketing of fresh or dried mangoes generated more than 21 millions USD [1]. Despite all these encouraging data, mango sector meets many phytosanitary difficulties that could compromise its productivity. Recently, a mango decline outbreak was observed in several production areas in Burkina Faso, affecting trees of all ages [2]. Prevalence of 25-100 % was reported with severity ranging from 25 % of affected plants in Kenedougou (South Sudanese zone), to 80 % in Nahouri (North Sudanese zone). These two main mangoes producing regions are located between the 10^th^ and 13^th^ parallel, have a tropical climate with contrasting rainy (4-5 months; 900-1100 mm annual precipitation) and dry seasons. Symptoms include dying back of twigs from the top, discolouration and the death of leaves, gummosis from branches, and browning of the vascular tissues when cross-sections were made in diseased mango twigs and main trunks. In the advanced stages of disease, the branches dry up one after the other, leading to partial or total decline of the trees a few months after first symptoms appeared [2].

Mango decline, or sudden death syndrome or mango dieback [3], is considered one of the most serious threats in all mango producing regions, leading to significant yield loss and low fruit quality of mango. Fungi from Botryosphaeriaceae family (Botyrosphaeriales: Dothideomycetes: Ascomycota) are frequently reported as causal agents of mango dieback, canker, and fruit rot in many countries [4,5]. Among them, several *Lasiodiplodia* species are now well recognized as dieback agents, including *Lasiodiplodia crassispora, L. egyptiacae, L. hormozganensis, L. iraniensis, L. pseudotheobromae* and *L. theobromae* [5,6,7,8]. They are latent endophytic fungi with a wide host ranges that cause disease when the host plant suffers stress, including drought and high temperatures [9,10]. *Lasiodiplodia* is described as a common soil-borne saprophyte, however, it is not clear how the fungus invade the plant, and locates inside vessels, the role of seed transmission, and pruning wounds in the epidemiology of the disease and if the fungus may be transmitted through the mango bark beetle, *Hypocryphalus mangiferae* (Stebbing). Factors such as poor agronomic management activities and climate warming may thus favor mango dieback caused by *Lasiodiplodia* spp.

The main objective of this work was to determine the diversity and pathogenicity of *Lasiodiplodia* species associated with mango dieback in Burkina Faso. A total of 47 isolates of *Lasiodiplodia* were used and identified through a combination of morphology and phylogenetic analysis based on the partial translation elongation factor 1-α (*tef1-α*) sequences and internal transcribed spacer (ITS) of the ribosomal DNA (rDNA) region. In addition, we evaluated the influence of macro environmental factors (climatic and edaphic features) on the disease intensity (incidence and severity) and distribution across the country.

## 2. Material and Methods

### 2.1. Sample collection

Samples were collected during field surveys carried out between March and November 2017 and in May 2018 in the six mango-producing provinces of Burkina Faso: Houet, Kénédougou, Comoé, Sanguié, Nahouri and Sissili (Figure 1) [2]. In 17 localities, 1-2 orchards were sampled. The samples consited of leaves, twigs or roots which were collected from randomly selected trees showing the characteristic symptoms of dieback. For each sample, the date, locality, mango variety and geographical coordinates, taken with GPS, were recorded. Samples were taken to the phytopathology laboratory of INERA (Institute of the Environment and Agricultural Research) Farako-bâ for pathogen isolation and the International molecular biology platform INERA/LMI PathoBios at Bobo Dioulasso for DNA analyses.

**Figure 1.**
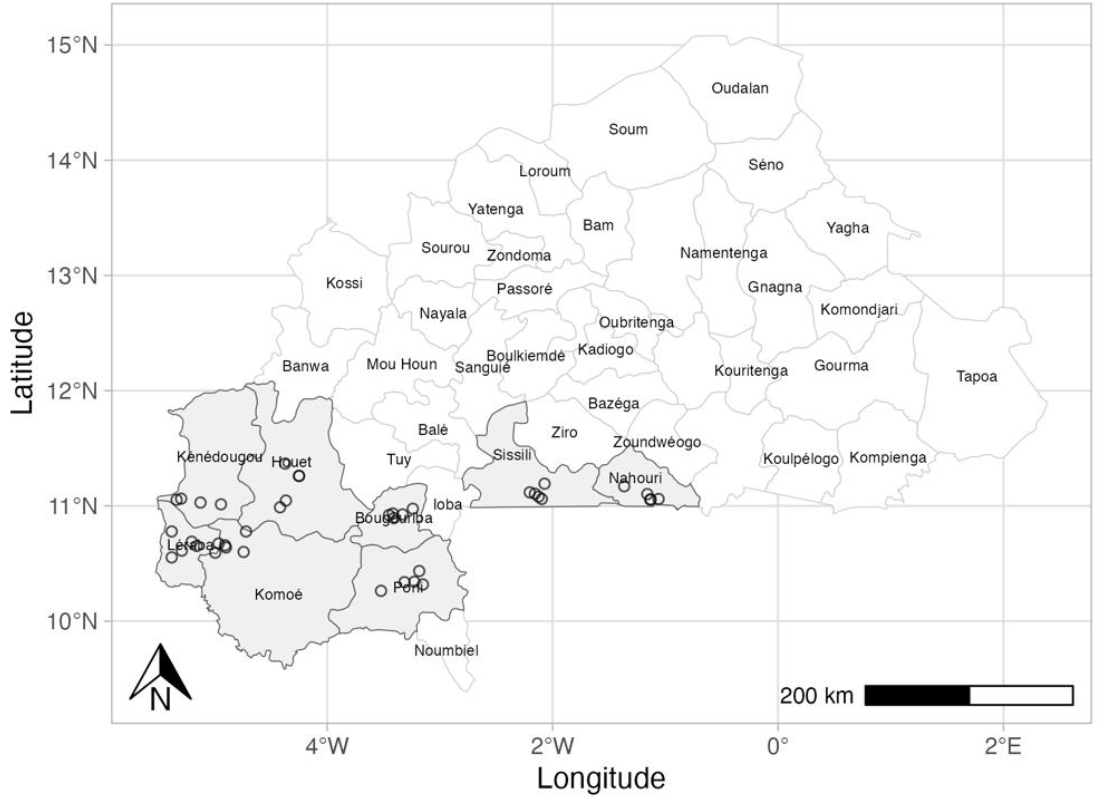
Map of Burkina Faso indicating the localities (empty circles) and provinces (grey fill) where field surveys and sampling were conducted.

### 2.2. Fungal isolation

Symptomatic samples were initially washed with tap water. Pieces from the edges of the symptomatic and symptomless tissues were cut and surface-sterilized with 70 % ethanol for 30 s followed by 1 % NaOCl for 1 min, rinsed in sterile distilled water for 30 s and incubated at room temperature (25-28°C) on a moistened blotting paper into Petri dishes. Eventually, hyphal tips of the fungal isolates were transferred onto fresh Potato Dextrose Agar (PDA) to obtain pure cultures.

### 2.3. Morphology of *Lasiodiplodia* spp isolates

The macro- and micro-morphological characteristics of twenty isolates were examined. Colony features and pigmentation were observed after seven days incubation on PDA, while the growth rate of each isolate was measured in triplicate after 72 h of incubation. Production of conidia were obtained by transfering a plug of 7-days mycelia culture on disinfested (as described above) mango leaves maintained on 1% water agar. The characteristics of conidia (shape, size, colour, longitudinal striations and presence or absence of septum) were observed after three weeks of incubation in a 12-h photoperiod of fungal cultures on PDA. Thirty conidia were measured for each isolate using a light microscope (Euromex, The Netherlands) with a micrometer at 40 X magnification.

### 2.4. DNA extraction and PCR

Approximately 20 mg of mycelium were scraped from the PDA surface, placed in a sterile 2 mL microcentrifuge tube, freezed with liquid nitrogen and grinded into a fine powder using a microcentrifuge tube pestle. Total DNA was extracted using the Plant Tissue protocol of the [11]. Quality and integrity of the DNAs were checked on an agarose electrophoresis gel. The ITS and *tef1-α* partial gene sequences of all fungal isolates were amplified by Polymerase Chain Reaction (PCR) from genomic DNA using universal fungal primers ITS4 (forward primer) and ITS5 (reverse primer) [12] or EF1-688-F and EF1-1251-R [13], respectively. Each PCR reaction was perfomed in a total reaction volume of 50µl including 10µl Hot FIREpol Blend Mastermix (Solis Biodyne, Tartu, Estonia), 2µl of 10µM each forward, and reverse primer, 4µl of genomic DNA (20ng/µl) and 32µl distilled water. All PCRs were carried out in a Thermo Hybaid PXE Thermal Cycler.

Amplification products were separated by 1% w/v agarose gel and stained with Ethidium bromide. PCR products were purified using the PCR Promega kit (Promega Corporation, WI, États-Unis) following the manufacturer’s instructions and sent to Genewiz company (www.genewiz.com/en-GB) for sequencing in both directions.

### 2.5. Phylogenetic analyses

Raw sequences were imported and analyzed in Geneious (version 8.1.8 created by Biomatters. available from http://www.geneious.com [14]. High quality reads (Phred >20) only were conserved and the forward and reverse sequences were aligned to generate a consensus sequence. Each consensus sequence was checked and manually edited when necessary. Genetic relatedness of isolates was assessed by multiple global alignement of ITS rDNA and *tef1-α* sequences using Geneious alignment. Unique ITS rDNA and *tef1-α* sequences obtained for a group of isolates were deposited at Genbank under accession numbers OQ054337-OQ054342 (ITS) and OP546359 - OP546364 (*tef1-α*) (Table 2).

The sequences obtained were compared with fungal sequences available in GenBank database using BlastN homology searches. For phylogenetic analyses, we retrieved the identical *Lasidiplodia* sequences from each locus (ITS rDNA and *tef1-α*) in GenBank (9 species), as well as from well characterized isolates (type collection) of 46 *Lasidiplodia* species listed and used in (Bezerra *et al*., 2021) ITS rDNA and *tef1-α* sequences of *Diplodia mutila* (strain CMW7060) were used as outgroup. Prior to phylogenetic analysis, ambiguous sequences at the start and end were deleted in order to optimize the alignments. Unique DNA sequences generated in this study were aligned along with DNA sequences retrieved from GenBank using the default settings of the ClustalW algorithm [15] in MEGA 11 [16]. Alignments were used to find the best nucleotide substitution model in MEGA 11 with the Bayesian Information Criteria (BIC) and Akaike Information Criteria (AIC). Individual data sets (ITS rDNA and *tef1-α*) were analyzed with the topological congruency test using Concaterpillar with P = 0.05 as cut off. Phylogenies were inferred in MEGA11 by using the Maximum Likelihood method as described in [17] The Nearest Neighbor Interchange (NNI) heuristic method was used for trees inference. The initial tree was automatically generated using the Maximum Parsimony method. No branch swap filter was selected. Bootstrap values for branch support were determined using 1,000 replications. All sites were included in the analysis

### 2.6. Pathogenicity tests

Pathogenicity tests were conducted for 12 randomly selected *Lasiodiplodia* isolates. The apical stem region of one year-old mango seedlings cv Amélie was surface-sterilized with 70% ethanol, and mechanical wounding was applied with sterilized scalpels. Plugs (5mm in diameter) of *Lasiodiplodia* spp mycelia were collected from PDA cultures of 7 days and applied on the plant wounded area. The inoculation area was then wrapped with cotton and parafilm as described in [8]. Control seedlings were treated with sterile PDA plugs. All mango seedlings were further maintained in a greenhouse with a photoperiod extended to 15 h under fluorescent lights at 28°C, and were examined for disease symptoms for 42 days. To satisfy Koch’s postulates, pieces of inoculated organs were removed from sites showing disease symptoms at 5 dpi, surface sterilized as mentioned above and plated on PDA. An isolate was considered pathogenic when it was able to cause disease symptoms and further isolated from the inoculated plant.

### 2.7. Incidence, severity and effect of climate and soil types on mango decline

On each sampling site, the disease incidence (proportion of infected trees) and severity were recorded as described in Dianda et al. [2]. Monthly summaries of meterological variables (mean, maximum and minimum temperature, dew point temperature, relative humidity, and precipitation) spanning five years prior to the survey (2013 to 2017) were obtained from the NASA POWER project via the nasapower R package version 4.0.9 [18]. The search was based on the geographic coordinates (latitude and longitude) for each of the 40 orchards located across eight provinces of Burkina Faso: Bougouriba, Comoé, Houet, Kénédougou, Léraba, Nahouri, Poni, Sanguié, and Sissili [2]. The soil type for each location was obtained from WorldClim Database [19].

The incidence and severity were plotted in a geographic map and their relationship was studied via linear regression modelling. Correlation analysis was used to determine the strength of the association between each pair of variables, considering all climatic variables (averaged across the five years) and mango decline incidence. All analysis were performed in R [20].

## 3. Results

### 3.1. Morphological characterization of *Lasiodiplodia* spp. isolates associated with mango decline

On PDA, colonies of *Lasiodiplodia* spp. isolates had initial white aerial mycelia that turned greenish-gray mycelium when they aged (Fig.1A). In general, isolates showed fast daily mycelial growth with the maximum radial growth of 8.5 cm obtained after 5 days. Conidia were obtained after 2-weeks culture on mango leaves maintained on water agar. Conidia shape was subovoid or ellipsoid, immature conidia were hyaline, one-celled and mature conidia were brownish to dark brown, two-celled with a thick wall and irregular longitudinal striations when aged (Fig.2 B&C). The size of conidia varied in average from 19.5 ± 1.4 µm (C17BF) to 22.7 ± 1.7 µm (H7BF) long and 10.5 ± 0.8 µm (C17BF) to 13.3 ± 1.2 µm wide (H31BF) (data not shown). All these morphological characteristics are typical of the genus *Lasiodiplodia* (Phillips *et al*., 2013).

**Figure 2.**
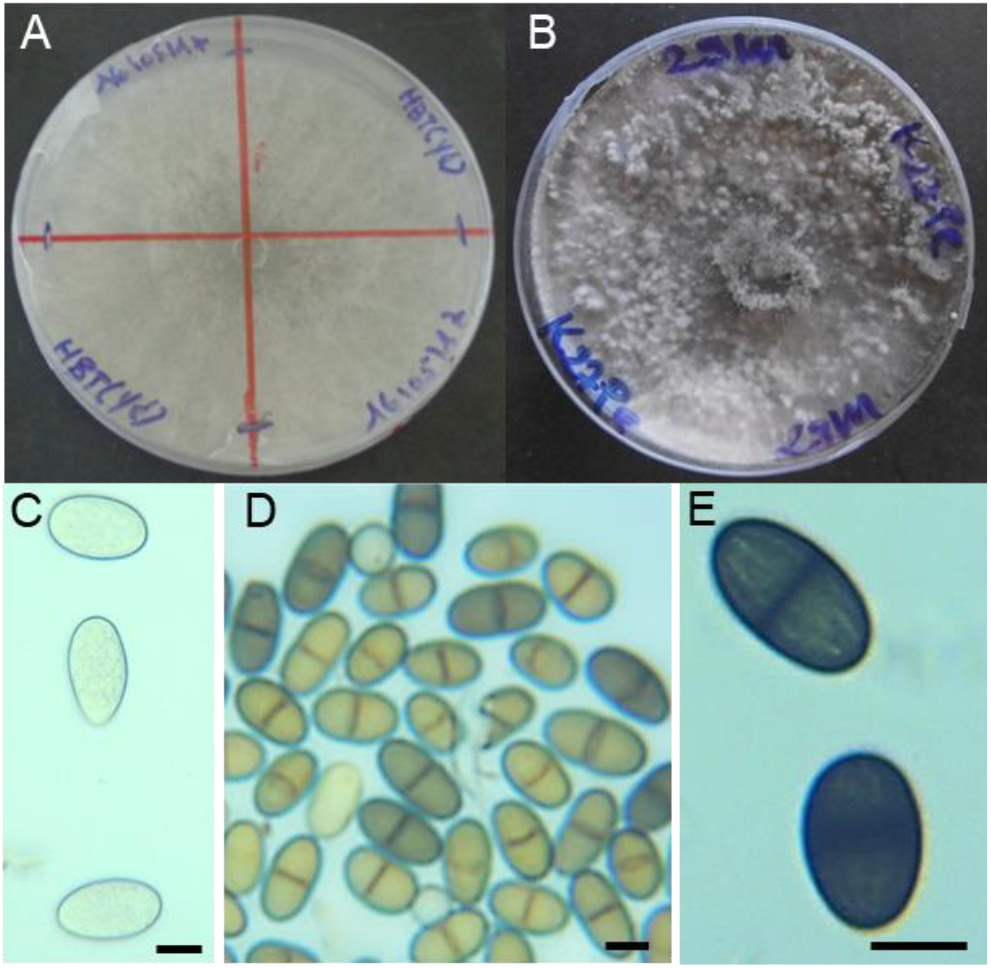
Morphological characteristics of *Lasiodiplodia* sp. growing on PDA plates. **A**: white aerial mycelia observed after 3 days; **B**: greenish-gray mycelium observed after 14 days; **C**: immature, hyaline conidia unicelled; D: brown conidia with septa; **C**: mature two-celled conidia with longitudinal striations. Bars: 10µm.

### 3.2. Molecular identification of *Lasiodiplodia* spp. isolates

We used the ITS rDNA and *tef1-α* partial gene to characterize 47 fungal isolates collected from diseased mango trees in six Burkinabé regions. ITS rDNA is widely used for taxonomic characterization in fungi, including the genus *Lasiodiplodia* but is not informative when used alone for discriminating between *Lasodiplodia* species. Combining ITS data with *tef1-α* data can help separating *Lasodiplodia* species [13]. We used the primer pair EF1-688F and EF1-1251R to amplify a *tef1-α* region that spans the first intron together with the part of the first and second exons [13]. Good quality sequences were obtained for ITS (average 538 bp, ranging from 429 to 549 bp) and for *tef1-α* (average 513, ranging from 455-548 bp) regions.

Alignment of ITS sequences showed that ITS sequences varied poorly among the 47 isolates, and were classified as A (and two 1-nt variants, A1 and A2), B (3-nt differences with A) and C (16-nt differences - including a 3-nt gap - with A). Regarding *tef1-α*, 6 distinct sequences (named A-F) were obtained among the 47 isolates, differing from 5 to 48 nt, from which 31 were parcimony informative. Final combined genotypes based on sequence data from the two loci enabled assignment of isolates to 6 ITS-*tef1-α* groups (I-VI) (Table 1). Sequences of representative isolates of the 6 genotypes found in Burkina Faso were deposited to GenBank (Table 2). β-tubulin (*tub2*) sequences were obtained for a subset of isolates from each of these genotypes but did not allow to separate more genotypes (data not shown).

**Table 1.** List of 47 *Lasiodiplodia* isolates obtained from symptomatic mango (*Mangifera indica*) trees in different regions in Burkina Faso and molecular classification based on ITS and *tef1-α* sequences obtained in this study.

| Isolate name | ITS | *tef1-α* | ITS-*tef1-α* genotype | Phylogenetically closest species | Province | Localities | Organ | Mango cultivar | GPS Coordinates |
| --- | --- | --- | --- | --- | --- | --- | --- | --- | --- |
| C2BF | A | B | I | *L. euphorbicola* | Comoé | Banfora | Twig | Mango vert | N10°65.762° W004°90.632° |
| C3BF | A2 | F | IV | *L. brasiliensis* | Comoé | Banfora | Leaf | Amélie | N10°65.762° W004°90.632° |
| C17BF | A | F | IV | *L. brasiliensis* | Comoé | Bérégadougou | Twig | Kent | N10°65.762° W004°90.632° |
| C19BF | A | B | I | *L. euphorbicola* | Comoé | Tingréla | Leaf | Amélie | N10°65.762° W004°90.632° |
| C20BF | B | E | III | *L. pseudotheobromae* | Comoé | Tingréla | Twig | Kents | N10°63.878°, W004°89.917° |
| H21BF | A | B | I | *L. euphorbicola* | Houet | Danfinso | Root | Kent | N11°260.17° W004°250.54° |
| H23BF | A | B | I | *L. euphorbicola* | Houet | Danfinso | Root | Brooks | N11°260.17° W004°250.54° |
| H24BF | A | B | I | *L. euphorbicola* | Houet | Darsalamy | Root | Brooks | N11°260.17° W004°250.54° |
| H22BF | A | A | II | *L. theobromae* | Houet | Darsalamy | Twig | Kent | N11°260.17° W004°250.54° |
| H25BF | B | E | III | *L. pseudotheobromae* | Houet | Darsalamy | Twig | Kent | N11°260.17° W004°250.54° |
| H30BF | A | B | I | *L. euphorbicola* | Houet | Peni | Root | Amelie | N11°155.75° W004°174.86° |
| H31BF | A | B | I | *L. euphorbicola* | Houet | Peni | Root | Amelie | N11°155.75° W004°174.86° |
| H37BF | A | B | I | *L. euphorbicola* | Houet | Peni | Leaf | Kent | N11°155.75° W004°174.86° |
| H7BF | A | B | I | *L. euphorbicola* | Houet | Yéguéresso | Root | Amelie | N11°046.080°W004°366.68° |
| H9BF | A | B | I | *L. euphorbicola* | Houet | Yéguéresso | Root | Amelie | N11°046.080°W004°366.68° |
| H15BF | A | B | I | *L. euphorbicola* | Houet | Yéguéresso | Root | Amelie | N10°98.781° W004°41.845' |
| K12BF | A1 | C | V | *L. caatinguensis* | KénéDougou | Dieri | Leaf | Lippens | N11°056.61' W005°196.12' |
| K7BF | A | A | II | *L. theobromae* | KénéDougou | Diéri | Root | Keitt | N11°056.61' W005°196.12' |
| K14BF | A | B | I | *L. euphorbicola* | KénéDougou | Koloko | Leaf | Brooks | N11°01.349'W004°56.581' |
| K16BF | A2 | F | IV | *L. brasiliensis* | KénéDougou | Koloko | Twig | Mango vert | N11°01.349'W004°56.581' |
| K6BF | A | B | I | *L. euphorbicola* | KénéDougou | Orodara | Root | Keitt | N11°056.61' W005°196.12' |
| K8BF | A | B | I | *L. euphorbicola* | KénéDougou | Orodara | Root | Valencia | N11°056.61' W005°196.12' |
| K9BF | A | C | V | *L. caatinguensis* | KénéDougou | Orodara | Twig | Amélie | N11°01.349'W004°56.581' |
| N1BF | A | B | I | *L. euphorbicola* | Nahouri | Pô | Root | Amelie | N11°0571.14' W001°078.39' |
| N8BF | A | B | I | *L. euphorbicola* | Nahouri | Pô | Root | Amelie | N11°102.33'W001°095.77' |
| N9BF | A | B | I | *L. euphorbicola* | Nahouri | Pô | Twig | Amelie | N11°102.33'W001°095.77' |
| N10BF | A | B | I | *L. euphorbicola* | Nahouri | Pô | Root | Amelie | N11°102.33'W001°095.77' |
| N11BF | A | B | I | *L. euphorbicola* | Nahouri | Pô | Root | Amelie | N11°102.33'W001°095.77' |
| N15BF | A | B | I | *L. euphorbicola* | Nahouri | Sango | Twig | Mango vert | N11°102.33'W001°095.77' |
| N14BF | B | E | III | *L. pseudotheobromae* | Nahouri | Sango | Twig | Amelie | N11°102.33'W001°095.77' |
| N12BF | C | D | VI | *L. crassispota* | Nahouri | Sango | Twig | Amelie | N11°102.33'W001°095.77' |
| N19BF | A | B | I | *L. euphorbicola* | Nahouri | Sarro | Twig | Amelie | N11°170.34'W001°219.06' |
| N17BF | A1 | B | I | *L. euphorbicola* | Nahouri | Sarro | Twig | Mango vert | N11°170.34'W001°219.06' |
| N18BF | A1 | C | V | *L. caatinguensis* | Nahouri | Sarro | Leaf | Amelie | N11°170.34'W001°219.06' |
| S6BF | A | B | I | *L. euphorbicola* | Sanguié | Secteur2 | Root | Amelie | N12°18.630' W002°25.194' |
| S10BF | A | B | I | *L. euphorbicola* | Sanguié | Secteur9 | Twig | Mango vert | N12°31.221' W002°46.463' |
| S15BF | A | B | I | *L. euphorbicola* | Sanguié | Secteur9 | Twig | Amelie | N12°31.221' W002°46.463' |
| I11BF | A | B | I | *L. euphorbicola* | Sissili | Gaoussa | Leaf | Amelie | N11°076.65° W002°133.50° |
| I10BF | A2 | A | II | *L. theobromae* | Sissili | Gaoussa | Twig | Amelie | N11°076.65° W002°133.50° |
| I2BF | A | B | I | *L. euphorbicola* | Sissili | Léo | Root | Amelie | N11°076.65° W002°133.50° |
| I6BF | A | B | I | *L. euphorbicola* | Sissili | Léo | Root | Mango vert | N11°076.65° W002°133.50° |
| I7BF | A | B | I | *L. euphorbicola* | Sissili | Léo | Root | Amelie | N11°076.65° W002°133.50° |
| I8BF | A | B | I | *L. euphorbicola* | Sissili | Léo | Root | Amelie | N11°076.65° W002°133.50° |
| I9BF | A | B | I | *L. euphorbicola* | Sissili | Léo | Twig | Amélie | N11°076.65° W002°133.50° |
| I13BF | A | B | I | *L. euphorbicola* | Sissili | Nadion | Twig | Amelie | N11°060.16° W002°0963.8° |
| I14BF | A | B | I | *L. euphorbicola* | Sissili | Nadion | Twig | Amelie | N11°060.16° W002°0963.8° |
| I15BF | A | B | I | *L. euphorbicola* | Sissili | Nadion | Twig | Amelie | N11°060.16° W002°0963.8° |

1 Acronyms of culture collections : **BF**= Burkina Faso; **L** = *Lasiodiplodia* ; **H** = Houet; **K** = Kénédougou; **N** = Nahouri; **C** = Comoé; **I** = Sissili; **S** 2 = Sanguie

**Table 2.** GenBank accession numbers of ITS and *tef1-α* sequences from representative isolates of *Lasiodiplodia sp*. genotypes found in Burkina Faso

| Isolate name | Phylogenetically closest species | Genbank accessions |  |
| --- | --- | --- | --- |
|  |  | ITS | *tefl-α* |
| H15BF | *L. euphorbicola* | OQ054341 | OP546363 |
| I10BF | *L. theobromae* | OQ054339 | OP546361 |
| H25BF | *L. pseudotheobromae* | OQ054340 | OP546362 |
| C3BF | *L. brasiliensis* | OQ054342 | OP546364 |
| K12BF | *L. caatinguensis* | OQ054338 | OP546360 |
| N12BF | *L. crassispota* | OQ054337 | OP546359 |

The genotype I was predominant with 34 of 47 Burkina isolates and present in all regions sampled. The remaining 13 isolates were separated into five genotypes (II to VI) disseminated in several regions (Table 1). Given the small number of isolates in each genotype, no inference about genetic differentiation based on geographic origin, organs or mango variety could be found.

Blast homology searches showed that the sequences obtained from isolates of diseased mango trees displayed high similarities with sequences of at least 8 *Lasidiplodia* species deposited in GenBank. Phylogenetic trees were inferred with individual datasets (ITS rDNA, *tef1-α*) for representative sequences of the 6 genotypes found in Burkina Faso (Table 2) and 45 *Lasiodiplodia* sequences retrieved from Genbank and [21] (suppl figures 1 and 2). The *Diplodia mutila* (strain CMW7060) sequences were used as outgroup. For the two datasets, the evolutionary history was inferred by using the Maximum Likelihood method and Kimura 2-parameter model [22] with a discrete Gamma distribution to model evolutionary rate differences among sites. There were a total of 460 positions in the final ITS dataset and 331 positions in the final *tef1-α* dataset. Reference isolates from Burkina Faso closely grouped with several *Lasodiplodia* species with high bootstrap values (68-100%) and, the *tef1-α* – based tree better assigned isolates to *Lasodiplodia* species than the ITS-based one (suppl figures 1 and 2). However, groupings were congruent in the two trees and Fig. 3 presents a *tef1-α* -based phylogenic tree constructed with a reduced number of *Lasodiplodia* species (9) that better matched the reference isolates from Burkina Faso. Genotype I reference isolate (H15BF) was found in a subclade with representative isolates of *L. parva* isolated from cacao (CBS356.59, Sri Lanka) and cassava (CBS456.78, Colombia) and *L. euphorbicola* CMM3609 isolated from Jatropha in Brazil. The closest relatives for the other *Lasiodiplodia* isolates from Burkina Faso were *L. theobromae* (I10BF, genotype II), *L. pseudotheobromae* and *L. lignicola* (H25BF, genotype III), *L brasiliense* and *L. viticola* (C3BF, genotype IV), or *L. caatingensis* (K12BF, genotype V). Isolate N12BF (genotype VI) was clearly separated from the others in a clade with *L. crassispora*.

**Figure 3.**
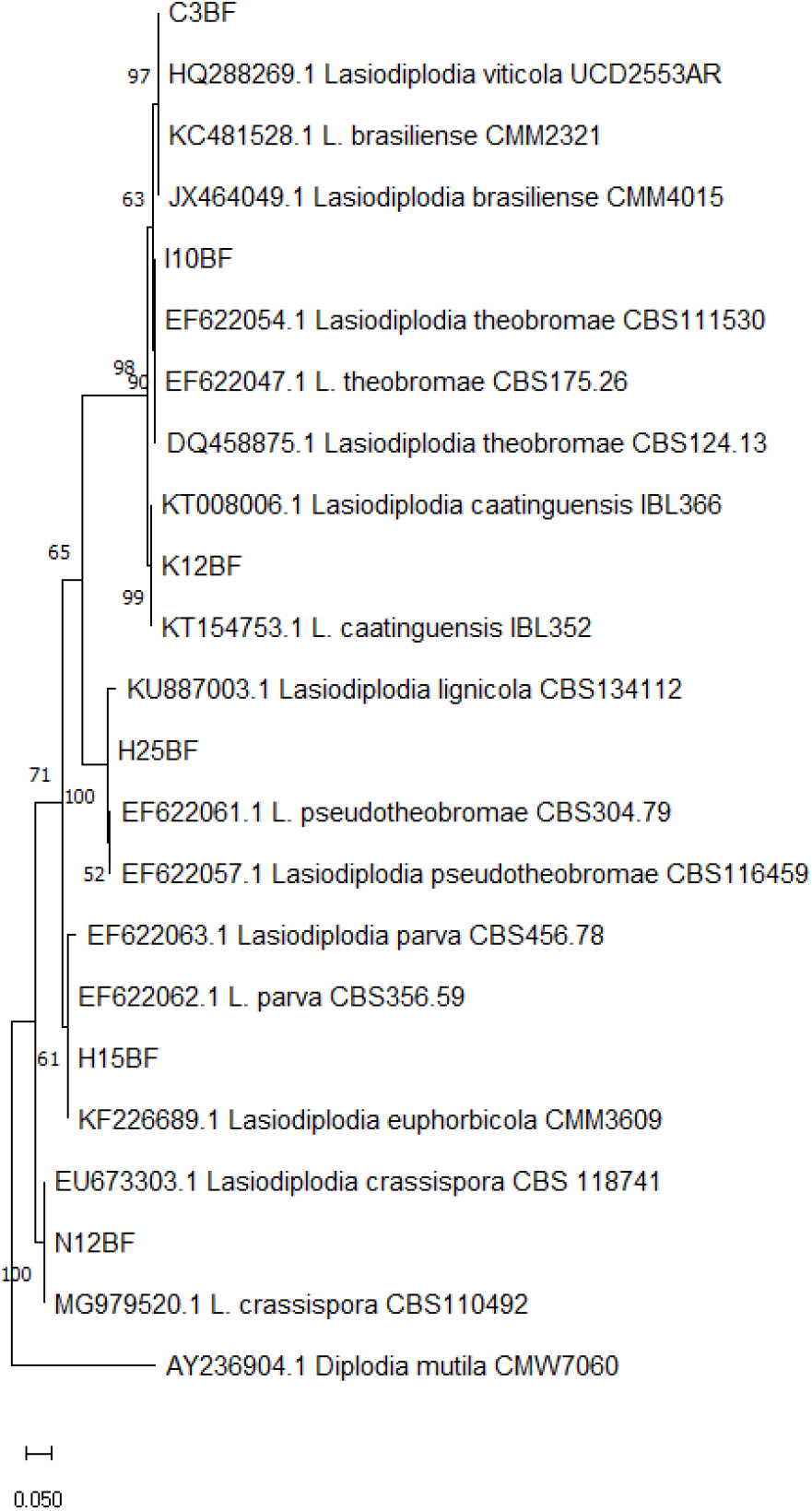
Maximum likelihood tree of the *tef1-α* dataset of 6 *Lasiodiplodia* spp. isolates from Burkina Faso and 16 *Lasiodiplodia* species isolates. The tree was generated based on the Tamura 3 parameter model using 1000 bootstraps. The tree with the highest log likelihood (-916.71) is shown. The percentage of trees in which the associated taxa clustered together is shown next to the branches. Initial tree(s) for the heuristic search were obtained automatically by applying the Maximum Parsimony method. A discrete Gamma distribution was used to model evolutionary rate differences among sites (5 categories (+G, parameter = 0.3237)). *Diplodia mutila* was used to root the tree. There were a total of 278 positions in the final dataset. Evolutionary analyses were conducted in MEGA11.

**Figure 3a:**
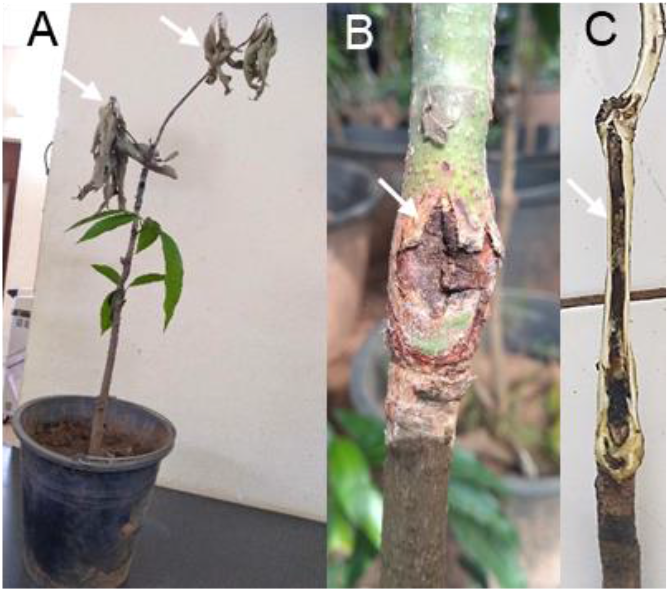
Dieback symptoms developed by mango seedlings *(M. indica* cv Amélie) 7 weeks after stem wound inoculation with *Lasiodiplodia* isolates from Burkina Faso. **A:** Wilting and death of the apical shoots with died leaves remaining attached (indicated by white arrows). **B**: necrosis and craking of the epidermal tissues around the inoculation site of inoculated stems (arrow), **C:** Necrosis and brown to black discolouration (arrow) of the vascular tissues extending upward and downward of the inoculation point.

This phylogenetic analysis based on ITS and *tef1-α* sequences variations showed that *Lasiodiplodia* isolates from diseased mango trees in Burkina Faso are genetically diversified and belong to different species. Even if a high number of isolates from different geographical regions grouped with reference *L. euphorbicola* and *L. parva* isolates, a genetic diversity was found in most of the regions sampled, with two species in Sissili, three in Comoé and Houet, four in Kénédougou and Nahouri (Table 1). At a small scale (locality/orchard), up to three species could be found in Darsalami (Houet) and Sango (Nahouri) localities.

### 3.3. Pathogenicity tests on mango seedlings

In parallel, a subset of 12 *Lasiodiplodia* isolates randomly chosen were tested for pathogenicity to mango seedlings. Inoculated seedlings developped typical dieback symptoms, including brown, necrotic bark lesions around the inoculation sites extending upwards and downwards, leading to wilting and drying of the apical leaves as well as the terminal leaves (Figure 3C). We also observed cracking of stem cortex or canker (Figure3 A). Under the outer cortex, necrotic xylem vessels and brown discolouration extended along the length of the stem (Figure 3 B). Control plants did not develop any symptoms.

All isolates tested were able to induce some characteristics dieback symptoms, and internal lesion length ranged from 2.76 ± 0,52 cm to 6.72 ± 0.61 cm (Table 3). *Lasiodiplodia* sp. were consistently re-isolated from the disease affected tissues; thus fulfilling Koch’s postulates that these detected symptoms were associated with inoculation of the fungus. No fungal isolate was obtained from the plant negative control.

**Table 3:** Number of plants displaying characteristic mango decline symptoms 7 weeks after stem wound inoculation with *Lasiodiplodia* isolates from Burkina Faso.

| Isolate | Phylogenetic group | Mean ± SD*<br>lesion size (cm) | % plants with<br>canker |
| --- | --- | --- | --- |
| N12BF | *L. crassisporea* | 2.76 ± 0.52 | 0 |
| H15BF | *L. euphorbicola* | 6.11 ± 0.46 | 75 |
| H23BF | *L. euphorbicola* | 4.53 ± 1.47 | 50 |
| H37BF | *L. euphorbicola* | 4.28 ± 0.65 | 50 |
| H9BF | *L. euphorbicola* | 3.22 ± 0.71 | 25 |
| K1BF | *L. euphorbicola* | 6.72 ± 0.61 | 50 |
| N10BF | *L. euphorbicola* | 4.10 ± 0.76 | 50 |
| N8BF | *L. euphorbicola* | 4.31 ± 0.62 | 50 |
| S10BF | *L. euphorbicola* | 5.61 ± 0.50 | 50 |
| S15BF | *L. euphorbicola* | 4.54 ± 0.63 | 50 |
| H25BF | *L. pseudotheobromae* | 5.76 ± 0.97 | 25 |
| N14BF | *L. pseudotheobromae* | 5.41 ± 0.93 | 25 |

\* SD: standard deviation; $n=4$ plants

### 3.4. Incidence, severity and climatic influence

The intensity of mango dieback (incidence and severity) followed a clear East-West (min to max) gradient. Increased disease intensity was reported in Nahouri and Sissili regions (North Sudanese climate – eastern regions) than in other Sub-Sudanese regions (Figure 5A,B). The incidence and severity were strongly associated and a second-order linear model fitted well the data, suggesting that severity could be predicted from incidence data (*R*^2^ = 0.82) (Figure 5C). Correlation analysis showed that longitude was significantly associated with disease incidence; the higher the longitude (eastern regions), the higher the disease. Mean annual temperature was positively associated (*r* = 0.7) with mango decline incidence, but inversely associated (*r* = - 0.64) with mean annual rainfall (Figure 6A). This suggests that higher disease intensity was observed in relatively drier and warmer regions. Soil types also seem to have influenced mango dieback with a generally higher incidence in Lithosols (Suppl. Fig. 3).

**Figure 5.**
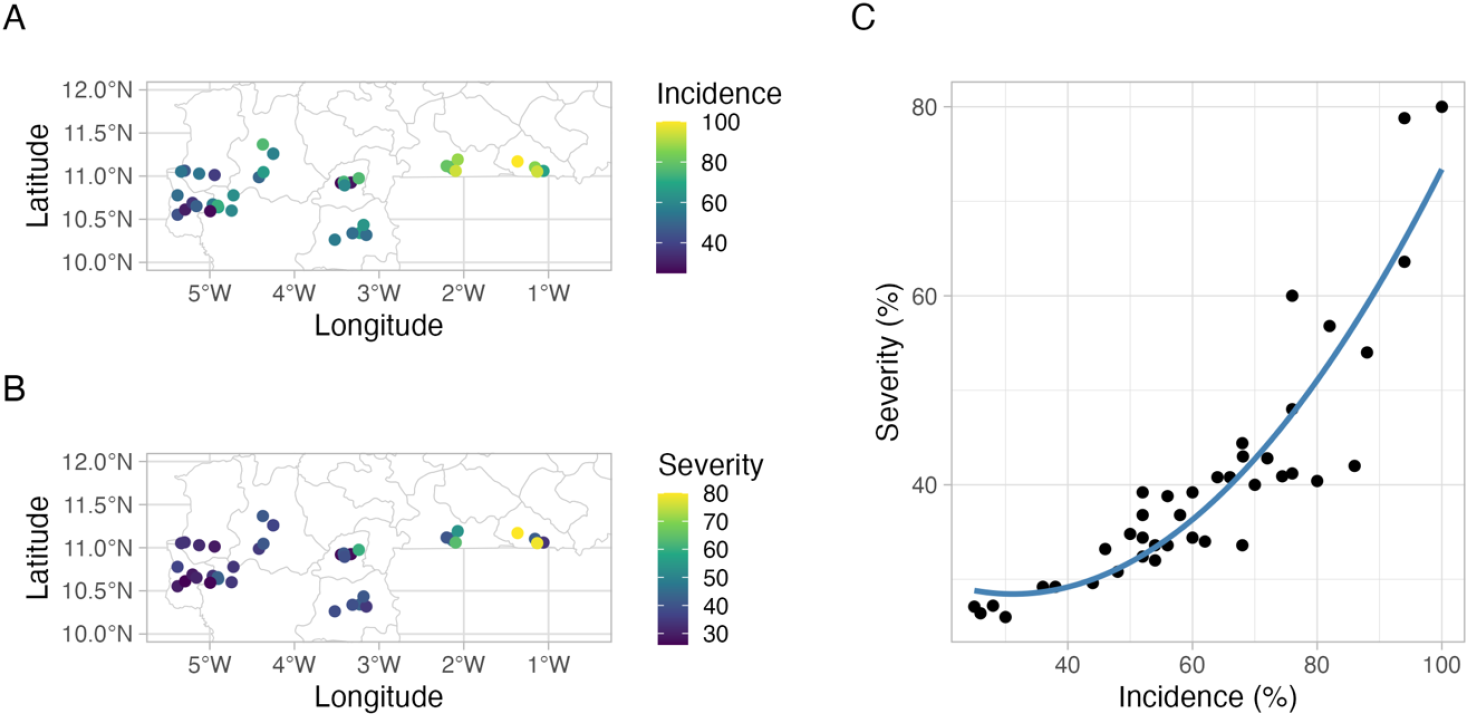
Geographical distribution of mango dieback incidence (A) and severity (B) in Burkina Faso and the linear relationship between these two variables (C). The solid blue line is the fit of a second-order linear regression model.

**Figure 6.**
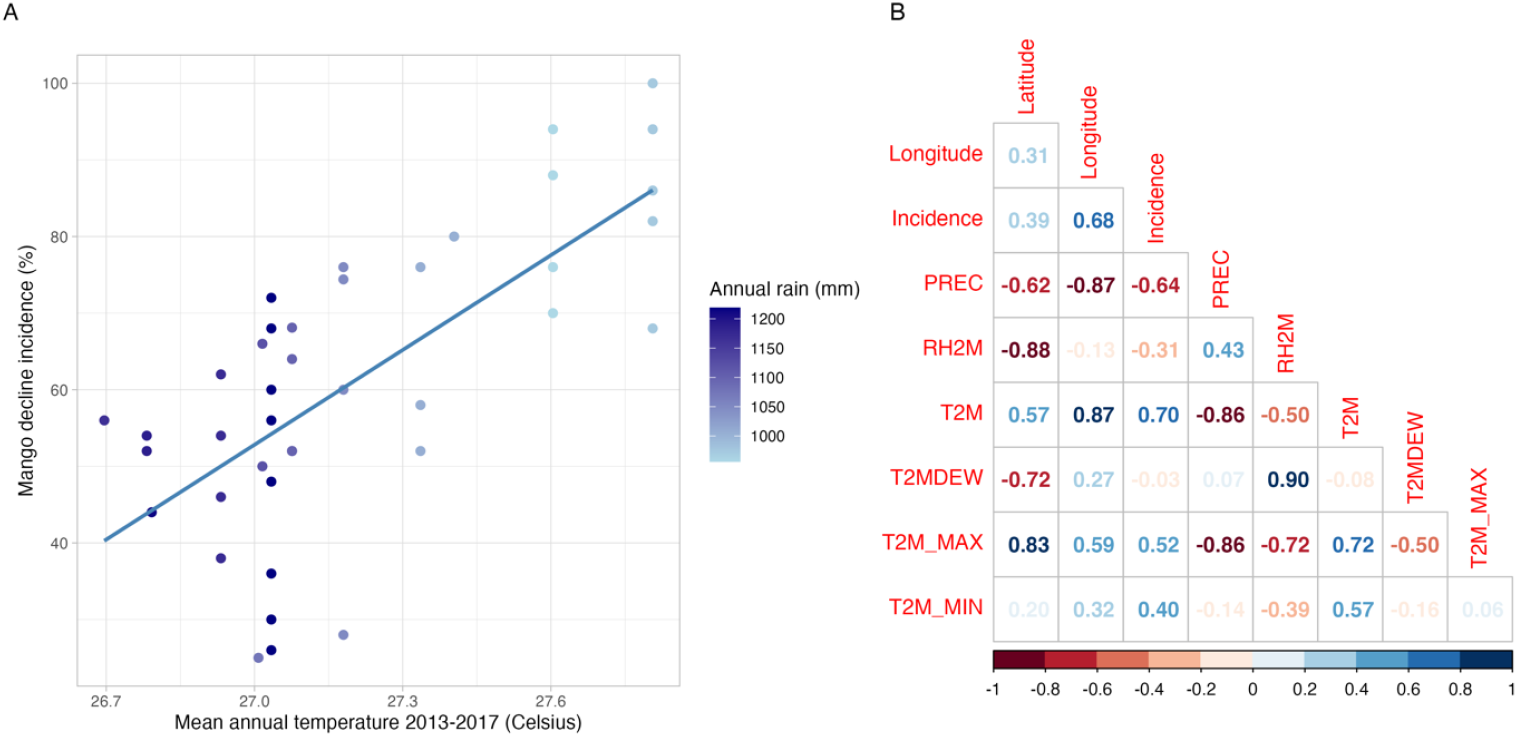
A. Relationship between mean annual temperature (dots) and mean annual rain (gradient color of the dots) averaged over five years and mango decline incidence. The blue solid line in A represents the fit of a first-order linear model. B. Correlation matrix for all pairs of variables.

## 4. Discussion

This study represents the first identification of *Lasiodiplodia* species associated with mango dieback in the main areas production of Burkina Faso, integrating morphology, pathology and molecular data. *Lasiodiplodia* isolated from symptomatic mango plants clustered with six species or species groups: *L. brasiliense* / *L. viticola, L. caatiguensis, L. lignicola* / *L. pseudotheobromae, L. theobromae, L. euphorbicola* / *L. parva*, and *L. crassispora*. This result indicates that dieback may be caused by a diverse group of *Lasiodiplodia* fungi dispersed in several mango-growing regions. *Lasiodiplodia* isolates tested in this study were pathogenic when inoculated on mango seedlings. These results on *Lasiodiplodia* species diversity are similar to those reported on mango dieback in other countries throughout the world [5,6,7,8]. Since the use of molecular analyses based on ITS and *tef1-α* sequences, several cryptic species were described resembling *L. theobromae* based on morphological characteristics [13]. Indeed, Phillips *et al*. [4] proposed a key to *Lasiodiplodia* species based on conidial dimensions and morphology of the paraphyses, however requiring an experienced technician and at least two weeks up to three months [23] of mycelia cultivation in specific conditions to induce sporulation. Here, we used molecular techniques to analyse isolates from diseased mango trees in Burkina Faso that possessed morphological features typical of the *Lasiodiplodia* genus, namely slowly maturing conidia with thick walls and longitudinal striations. Isolates collected in mango orchards of Burkina Faso and tested on mango seedlings (variety Amelie) were pathogenic, producing various symptoms characteristics of mango dieback. These data support findings of previous studies showing that inoculation of mango plants with *Lasiodiplodia* species can manifest various external and internal symptoms such as bark necrosis, vascular discolouration, defoliation, apical dieback and gummosis [8,24,25,26,27]. *Lasiodiplodia* species are frequently reported as the most aggressive pathogens of mango among the wide diversity of *Botryosphaeriaceae* found on this host. The different *Lasiodiplodia* species were distributed in all regions sampled, with the predominance of a single species (*L. euphorbicola*) throughout the country. However, up to three species could also be found in a single locality (Table1), indicating that dieback maybe caused by different *Lasiodiplodia* isolates even at a small geographic scale. In addition to mango, phylogenetic studies have identified a diversity of *Lasiodiplodia* species associated to dieback of other fruit species. For example, eight distinct *Lasiodiplodia* species were identified on table grape dieback in Brazil [28]. Four species of *Lasiodiplodia* were associated with *M indica, Citrus reticulata, C. sinensis, Ficus carica, Prunus persica, Prunus armeniaca* and *Pyrus communis* trees dieback [29]. Taking adavantge of new sequencing methods could be useful for accurately detecting and analyzing the microbiota associated to diseased orchards. For instance, nanopore metagenomics quickly identified *L. theobromae* among other fungi from root-soil samples and infected plant tissues collected from diseased mango trees in Egypt [30]. Alternatively, genotyping by sequencing (GBS) may be useful for investigating the pathogen molecular genetic diversity in several orchards through genome-wide SNP discovery, enabling further genetic population studies for review see [30]. Knowledge about *Lasiodiplodia* spore production and propagation in orchards is also limited. Such investigations could be very informative for epidemiological studies of the disease progression. Understanding how *Lasiodiplodia* species and genotypes spread to new trees in orchards or new aereas could help control the pathogen.

In this study we examined for the first time bioclimatic factors favouring mango dieback risk at large spatial scales. We could establish that a higher prevalence was associated to higher annual mean temperature and lower rainfall in Burkina Faso when disease data were recorded (2018). Annual mean temperature in the localities sampled varied between 26 and 28.5°C, and annual rainfall varied between 900 and 1100 mm. It is well-known that heat stress and drougth can affect plant interactions with microbiota, including *Botryosphaeraceae* fungi [9]. Transcriptomic analyses showed that *Lasiodiplodia* do not turn more aggressive with increasing temperature, rather it seems that the plant host defence weakens and becomes more susceptible to the fungal endophyte [32]. In addition, climatic conditions may affect conditions that endorse pycnidia production and spore release, and favor infection. Orchards management methods for reducing the stress should take into account the seasonal susceptibility of pruning wounds in accordance with spore production. Searching for mango varieties more tolerant to stress induced by climate changes could help reducing the impact of dieback.

## 5. Conclusions

The study consisted of characterizing *Lasiodiplodia* spp. isolated from organs of mango suffering dieback in Burkina Faso. Morphological characteristics combined with the molecular tests permited to identify *L. brasiliensis, L. caatingensis, L. crassispora, L. euphorbicola, L. pseudotheobromae* and *L. theobromae* species associated to diseased mango trees. Climate characteristics do have an impact on the disease development. Additional studies are therefore required to prevent occurrence of this disease in mango orchards, and to develop a reliable and sustainable preventive control strategy.

## 6. Acknowledgements

The meterological data gathered using the Nasa power R package were obtained from the NASA Langley Research Center POWER Project funded through the NASA Earth Science Directorate Applied Science Program.. This work benefited from the facilities of the “International joint Laboratory (LMI) INERA/IRD PathoBios: Observatory of plant pathogens in West Africa: biodiversity and biosafety” (www.pathobios.com; twitter.com/PathoBios).

## Supplementary Figures

**Supplemental Figure 1:**
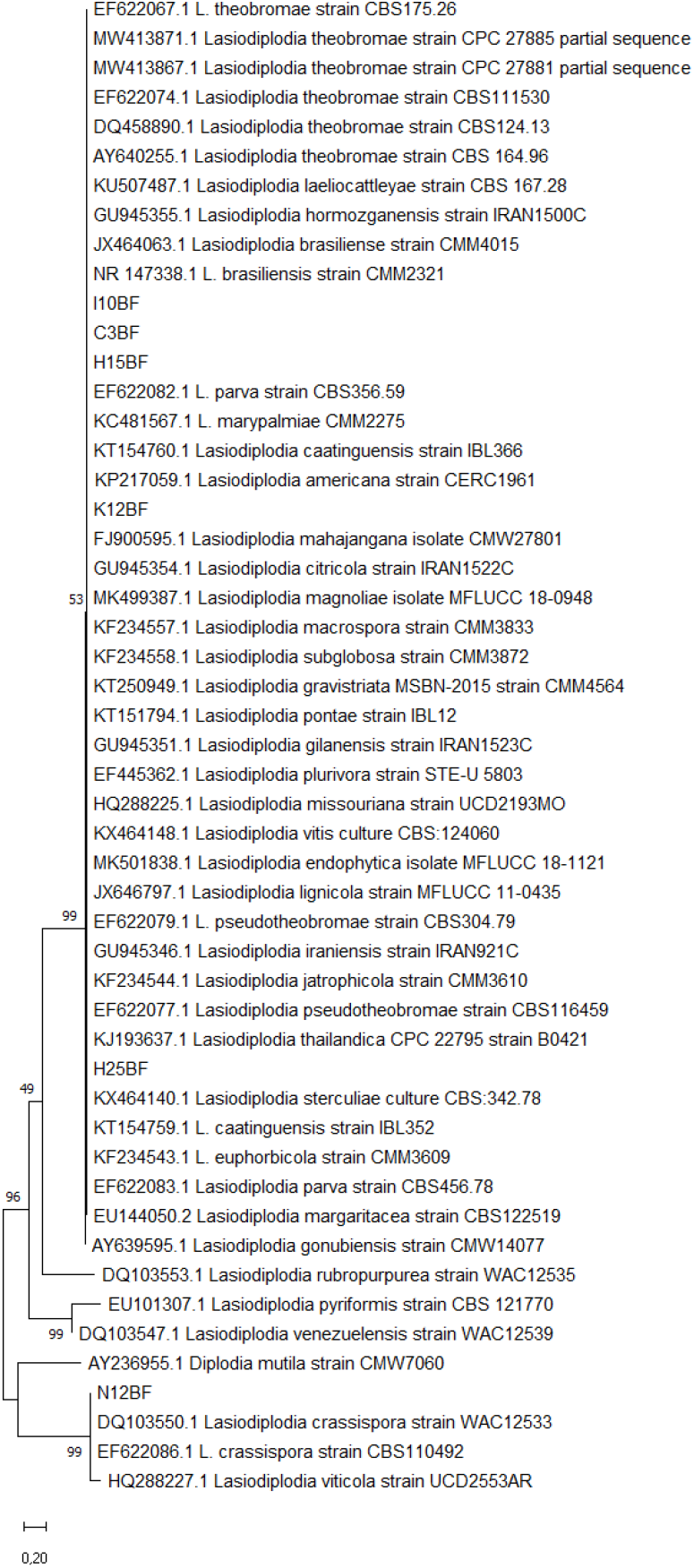
Maximum likelihood tree of the ITS dataset of 6 *Lasiodiplodia* spp. isolates from Burkina Faso and 44 *Lasiodiplodia* species isolates. The tree was generated based on the Tamura 3 parameter model using 1000 bootstraps. The tree with the highest log likelihood (-3539,90) is shown. The percentage of trees in which the associated taxa clustered together is shown next to the branches. Initial tree(s) for the heuristic search were obtained automatically by applying the Maximum Parsimony method. A discrete Gamma distribution was used to model evolutionary rate differences among sites (5 categories (+G, parameter = 200,0000)). *Diplodia mutila* was used to root the tree. There were a total of 460 positions in the final dataset. Evolutionary analyses were conducted in MEGA11.

**Supplemental Figure 2:**
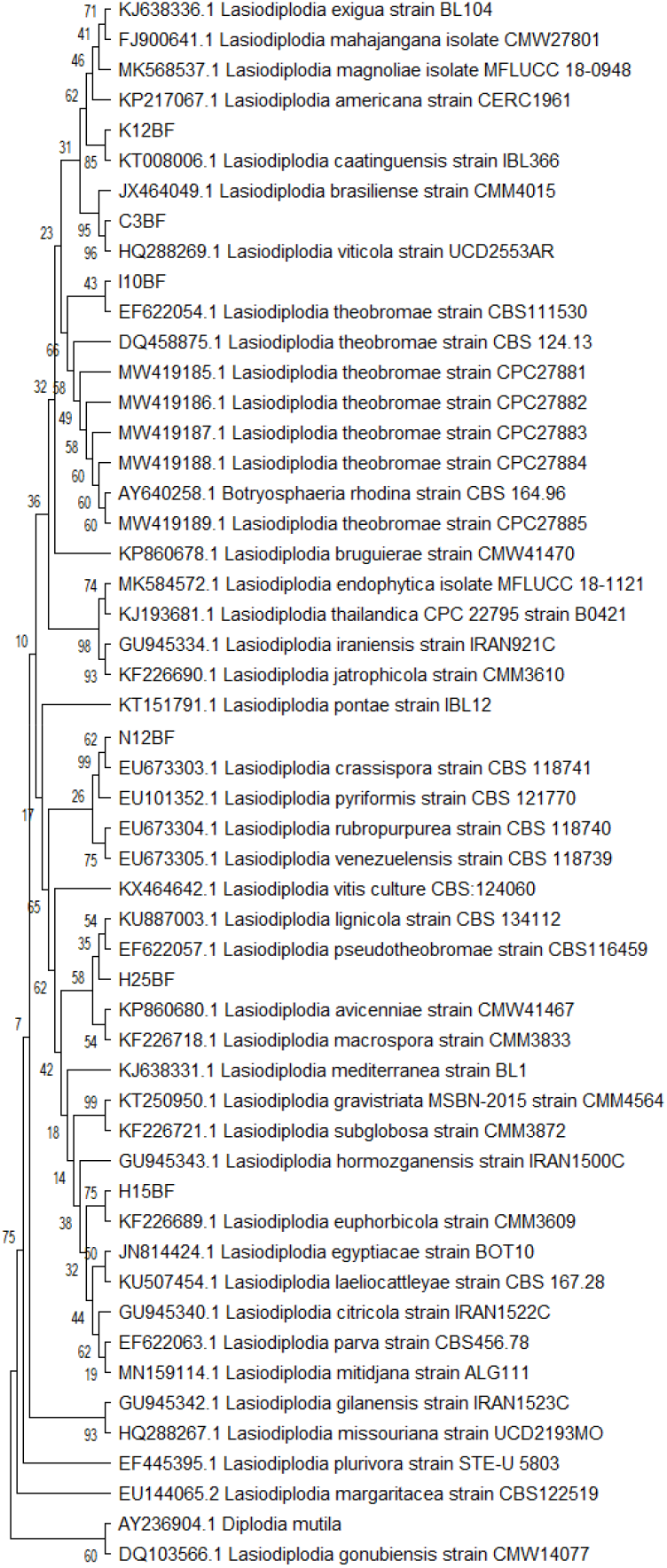
Maximum likelihood tree of the *tef1-α* dataset of 6 Lasiodiplodia spp. isolates from Burkina Faso and 45 Lasiodiplodia species isolates. The tree was generated based on the Tamura 3 parameter model using 1000 bootstraps. The tree with the highest log likelihood (-2154,11) is shown. The percentage of trees in which the associated taxa clustered together is shown next to the branches. Initial tree(s) for the heuristic search were obtained automatically by applying the Maximum Parsimony method. A discrete Gamma distribution was used to model evolutionary rate differences among sites (5 categories (+*G*, parameter = 0,4300)). *Diplodia mutila* was used to root the tree. There were a total of 331 positions in the final dataset. Evolutionary analyses were conducted in MEGA11.

**Supplemental Figure 3.**
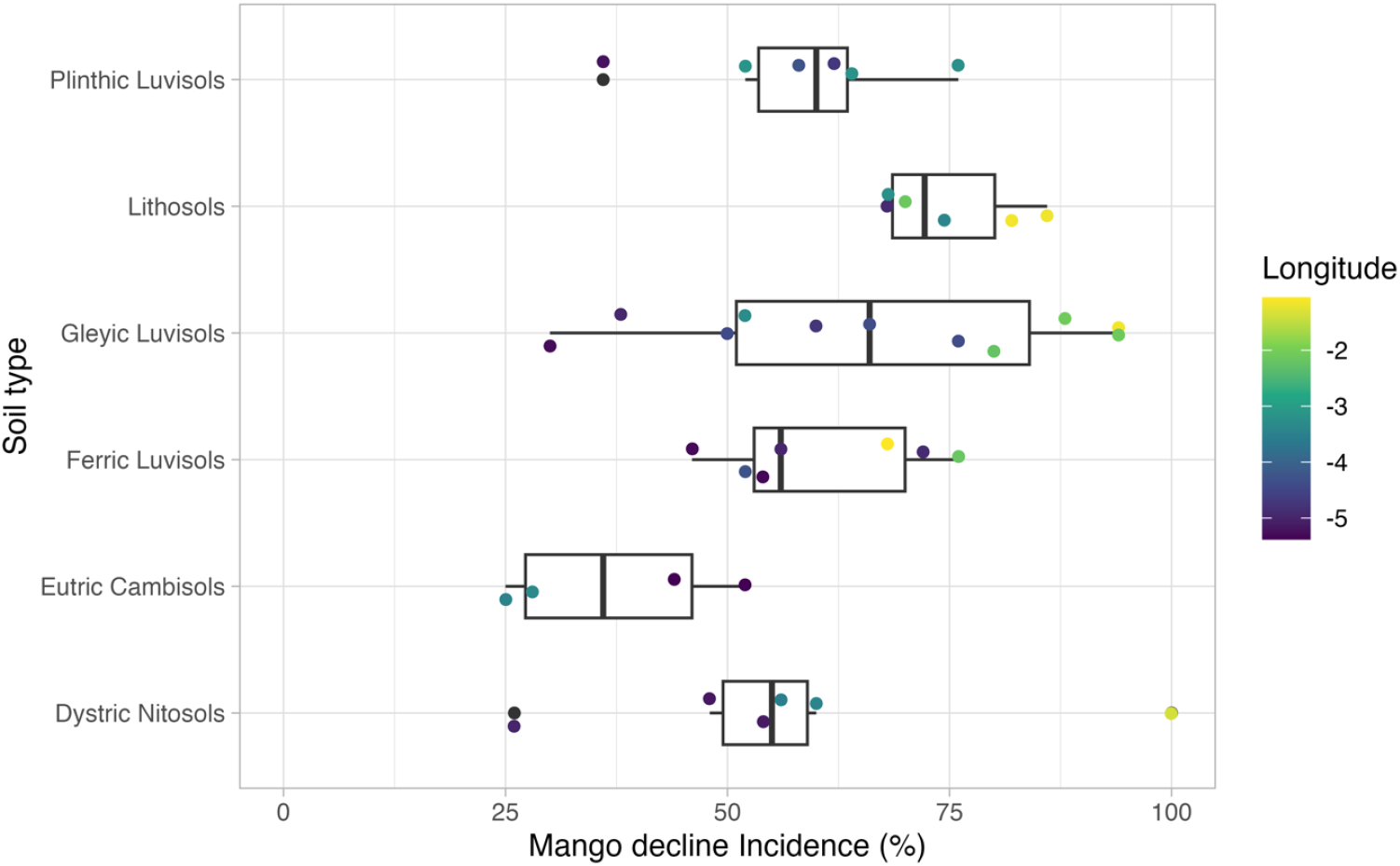
Box plot for the distribution of mango decling severity conditioned to soil types and longitude in Burkina Faso.

## References

[1] APROMAB, 2022. Lancement de la campagne mangue 2022 sous le thème: Réduire la pression parasitaire pour la campagne mangue 2022. Report 16 pages.

[2] Dianda, Z.O., Wonni, I., Zombré, C., Traoré, O., Sérémé, D., Boro, F., Ouédraogo, I., Ouédraogo, S.L., Sankara, P., 2018. Prévalence du dessèchement du manguier et evaluation de la fréquence des champignons associés à la maladie au Burkina Faso. J. Appl. Biosci. 126, 12686–12699. https://doi.org/10.4314/jab.v126i1.6

[3] Khaskheli, M.I., Jiskani, M.M., Soomro, M.H., Talpur, M.A., Poussio, G.B., 2011. Prevalence of Mango Sudden Decline / Death Syndrome (Msds) on Various Varieties At the Orchards of Different Age in the Vicinity of Tando Qaiser, Hyderabad, Sindh. Pak. J. Agri., Agril. Engg., Vet. Sci., 27, 160–167.

[4] Phillips, A. J. L., Alves, A., Abdollahzadeh, J., Slippers, B., Wingfield, M. J., Groenewald, J. Z., & Crous, P. W. (2013). The Botryosphaeriaceae: Genera and species known from culture. Studies in Mycology, 76, 51–167. https://doi.org/10.3114/sim0021

[5] Trakunyingcharoen, T., Cheewangkoon, R., To-Anun, C., Crous, P.W., Van Niekerk, J.M., Lombard, L., 2014. Botryosphaeriaceae associated with diseases of mango (Mangifera indica). Australas. Plant Pathol. 4, 425–438. https://doi.org/https://doi.org/10.1007/s13313-014-0284-9

[6] Ismail, A.M., Cirvilleri, G., Polizzi, G., Crous, P.W., Groenewald, J.Z., Lombard, L., 2012. Lasiodiplodia species associated with dieback disease of mango (Mangifera indica) in Egypt. Australas. Plant Pathol. 41, 649–660. https://doi.org/10.1007/s13313-012-0163-1

[7] Marques, W.M., Lima, B.N., Morais Jr, A. de M., Barbosa, G.A.M., Souza, O.B., Michereff, J.S., Phillips, J.L.A., Câmara, P.S.M., 2013. Species of Lasiodiplodia associated with mango in Brazil. Fungal Divers. 1947, 181–193. https://doi.org/10.1007/s13225-013-0231-z

[8] Rodríguez-Gálvez, E., Guerrero, P., Barradas, C., Crous, P.W., Alves, A., 2017. Phylogeny and pathogenicity of Lasiodiplodia species associated with dieback of mango in Peru. Fungal Biol. 121, 452–465. https://doi.org/10.1016/j.funbio.2016.06.004

[9] Slippers, B., Wingfield, M.J., 2007. Botryosphaeriaceae as endophytes and latent pathogens of woody plants: diversity, ecology and impact. Fungal Biol. 21, 90–106. https://doi.org/10.1016/j.fbr.2007.06.002

[10] Salvatore, M. M., Andolfi, A., Nicoletti, R., 2020. The thin line between pathogenicity and endophytism: The case of Lasiodiplodia theobromae. Agriculture, 10, 1–22. https://doi.org/10.3390/agriculture10100488

[11] Pinho, D.B., Firmino, A.L., Fereira, W.G., Ferreira-junior, L. Olinto, 2012. An efficient protocol for DNA extraction from Meliolales and the description of Meliola centellae sp. nov. Mycotaxon 122, 333–345. https://doi.org/10.5248/122.333

[12] White, T.J., Bruns, T., Lee, S., Taylor, J., 1990. Amplification and direct sequencing of fungal ribosomal RNA genes for phylogenetics. In: Innis M.A, Gelfand D.H, Sninsky J.J, White T.J, (eds.) (pp.). PCR Protocols : A Guide to Methods and Applications, Academic Press, New York, 315-322. http://dx.doi.org/10.1016/B978-0-12-372180-8.50042-1

[13] Alves, A., Crous, P.W., Correia, A., Phillips, J.L.A., 2008. Morphological and molecular data reveal cryptic speciation in Lasiodiplodia theobromae. Fungal Divers. 28, 1–13.

[14] Kearse, M., Moir, R., Wilson, A., Stones-Havas, S., Cheung, M., Sturrock, S., 2012. Geneious Basic: an integrated and extendable desktop software platform for the organization and analysis of sequence data. Bioinformatics 28, 1647e1649.

[15] Thompson, J.D., Gibson, T.J., Plewniak, F., Jeanmougin, F., Higgins, D.G., 1997. The CLUSTAL_X windows interface: flexible strategies for multiple sequence alignment aided by quality analysis tools. Nucleic Acids Res. 25, 4876–4882.

[16] Tamura, K., Stecher, G., Kumar, S., 2021. MEGA 11: Molecular Evolutionary Genetics Analysis Version 11. Mol. Biol. Evol. https://doi.org/https://doi.org/10.1093/molbev/msab120.

[17] Flor, N.C., Wright, A.F., Huguet-Tapia, J., Harmon, P.F., Liberti, D., 2022. Identification of Fungi in the Botryosphaeriaceae Family Associated with Stem Blight of Vaccinium spp. in the Southeastern United States. Fungal Biol. 5, 342–355. https://doi.org/https://doi.org/10.1016/j.funbio.2022.03.004

[18] Sparks, (2018). nasapower: A NASA POWER Global Meteorology, Surface Solar Energy and Climatology Data Client for R. Journal of Open Source Software, 3(30), 1035. https://doi.org/10.21105/joss.01035

[19] Fick, S.E., Hijmans, R.J., 2017. WorldClim 2 : new 1km spatial resolution climate surfaces for global land areas. Int. J. Climatol. 12, 4302–4315.

[20] R Core Team (2021). R: A language and environment for statistical computing. R Foundation for Statistical Computing, Vienna, Austria. URL https://www.R-project.org/.

[21] Bezerra, J.D.P., Crous, P.W., Aiello, D., Gullino, M.L., Polizzi, G., Guarnaccia, V., 2021. Diversity and Pathogenicity of Botryosphaeriaceae Species Associated with Symptomatic Citrus Plants in Europe. Plants 10, 492. https://doi.org/https://doi.org/10.3390/plants10030492

[22] Kimura M. (1980). A simple method for estimating evolutionary rate of base substitutions through comparative studies of nucleotide sequences. J. Mol. Evol. 16:111–120.

[23] Xia, G., Manawasinghe, I.S., Phillips, A.J.L., You, C., Jayawardena, R.S., Luo, M., Hyde, K.D., 2022. Lasiodiplodia fici sp. nov ., Causing Leaf Spot on Ficus altissima in China. Pathogens 11, 1–13. https://doi.org/https://doi.org/10.3390/pathogens11080840

[24] Ramos, L.J., Lara, S.P., McMillan Jr, R.T., Narayanan, R.K., 1991. Tip dieback of mango (Mangifera indica) caused by Botryosphaeria ribis. Plant Dis. 75, 315–318.

[25] Ploetz, R.C, Benscher, D., Vazquez, A., Colls, A., Nagel, J., Schaffer, B., 1996. A reexamination of mango decline in Florida. Plant Dis. 80, 664–668.

[26] Al-Jabri, M.K., Al-Shaili, M., Al-Hashmi, M., Nasehi, A., Al-Mahmooli, I.H., Al-Sadi, A.M., 2017. Characterization and evaluation of fungicide resistance among Lasiodiplodia theobromae isolates associated with mango dieback in Oman. J. Plant Pathol. 99, 753–759. https://doi.org/10.4454/jpp.v99i3.3959

[27] Saeed, E.E., Sham, A., AbuZarqa, A., Khawla, A.A.S., Tahra, S.A.N., Rabah, I., Khaled, E.-T., AbuQamar, S.F., 2017. Detection and Management of Mango Dieback Disease in the United Arab Emirates. Int. J. Mol. Sci. 18, 2086. https://doi.org/10.3390/ijms18102086

[28] Correia, K.C., Silva, M.A., Jr, Armengol, M.A.D.M.J., Phillips, A.J.L., Camara, M.P.S., Michereff, S.J., 2016. Phylogeny, distribution and pathogenicity of Lasiodiplodia species associated with dieback of table grape in the main Brazilian exporting region. Plant Pathol. 65, 92–103. https://doi.org/10.1111/ppa.12388

[29] El-Ganainy, S.M., Ismail, A.M., Iqbal, Z., Elshewy, E.S., Alhudaib, K.A., Almaghasla, M.I., Magistà, D., 2022. Diversity among Lasiodiplodia Species Causing Dieback, Root Rot and Leaf Spot on Fruit Trees in Egypt, and a Description of Lasiodiplodia newvalleyensis sp. nov. J. Fungi 8, 1203.

[30] Hassan, A.H., Bebawy, A.S., Saad, M.T., Mosaad, G.S., Saad, B.T., Eltayeb, W.N., Aboshanab, K.M., 2022. Metagenomic nanopore sequencing versus conventional diagnosis for identification of the dieback pathogens of mango trees. Biotechniques 6, 261–272.

[31] Vaux, F., Dutoit, L., Fraser, C.I., Waters, J.M., 2022. Genotyping‐by‐sequencing for biogeography. J. Biogeogr. 262–281. https://doi.org/https://doi.org/10.1111/jbi.14516

[32] Félix, C., Meneses, R., Gonçalves, M.F.M., Tilleman, L., Duarte, A.S., Jorrín-Novo, J. V, Van de Peer, Y., Deforce, D., Van Nieuwerburgh, F., Esteves, A.C., Alves, A., 2019. A multi-omics analysis of the grapevine pathogen Lasiodiplodia theobromae reveals that temperature affects the expression of virulence- and pathogenicity-related genes. Sci. Rep. 1, 1–12. https://doi.org/https://doi.org/10.1038/s41598-019-49551-w

